# Sensation and expectation are embedded in mouse motor cortical activity

**DOI:** 10.1101/2023.09.13.557633

**Authors:** Brooke E. Holey, David M. Schneider

## Abstract

During behavior, the motor cortex sends copies of motor-related signals to sensory cortices. It remains unclear whether these corollary discharge signals strictly encode movement or whether they also encode sensory experience and expectation. Here, we combine closed-loop behavior with large-scale physiology, projection-pattern specific recordings, and circuit perturbations to show that neurons in mouse secondary motor cortex (M2) encode sensation and are influenced by expectation. When a movement unexpectedly produces a sound, M2 becomes dominated by sound-evoked activity. Sound responses in M2 are inherited partially from the auditory cortex and are routed back to the auditory cortex, providing a path for the dynamic exchange of sensory-motor information during behavior. When the acoustic consequences of a movement become predictable, M2 responses to self-generated sounds are selectively gated off. These changes in single-cell responses are reflected in population dynamics, which are influenced by both sensation and expectation. Together, these findings reveal the rich embedding of sensory and expectation signals in motor cortical activity.

## Introduction

Behaviors often have predictable sensory consequences. For example, when shutting a car door, we expect to hear a stereotypical “thump” at a particular phase of our arm movement. Humans, monkeys, mice, and other animals can learn to predict what sound a movement will produce and when that sound will occur (Curio et al., 2000; Eliades and Wang, 2008; Schneider et al., 2014; Audette et al., 2022). At a behavioral level, animals use predictions about self-generated sounds to facilitate perception, to detect errors during sensory-motor execution, and to use errors to update subsequent behavior (Schneider et al., 2018; Eliades and Wang, 2008; Hickok et al., 2011; Houde and Jordan, 1998).

The ability to anticipate the sensory consequences of one’s actions is mediated through long-range connections between sensory and motor regions of the brain. Motor cortical neurons that are involved in executing and controlling behavior send copies of motor-related commands – termed corollary discharge – to sensory cortical areas, where they are integrated with incoming sensory signals (Peters et al., 2014; Park et al., 2022; Schneider et al., 2014; Crapse and Sommer, 2008; Nelson et al., 2013; Leinweber et al., 2017). In the auditory cortex, inputs from the secondary motor cortex (M2) target excitatory and inhibitory neurons in both deep and superficial layers, drive feedforward inhibition, and suppress neural responses to sounds during behaviors including locomotion, vocalizing, forelimb movements, grooming, and fidgeting (Nelson et al., 2013; Schneider et al., 2014; Audette et al., 2022; Morandell et al., 2023).

In addition to their role in the broad modulation of auditory cortex activity during movement, corollary discharge signals are also implicated in learning to anticipate the specific sensations an action is likely to produce, referred to as an internal model (Schneider and Mooney 2015; Schneider and Mooney 2018). During sound-generating behaviors, the auditory cortex simultaneously receives ascending acoustic information arriving from the periphery and top-down input about behavior (e.g. from M2). In the auditory cortex, experience with a sound-generating behavior results in the selective suppression of a movement’s expected consequences but weaker or no suppression when the same movement makes an unexpected sound (Schneider et al., 2018; Rummel et al., 2016; Audette et al., 2022; Audette et al., 2023).

Most models of motor-sensory learning presume that corollary discharge signals provide a stable representation of an animal’s current behavioral state so as to temporally coincide with and modulate self-made sensory input (Sperry, 1950, Straka et al., 2018). However, M2 activity does not strictly encode action. For example, M2 activity is modulated by reward and behavioral context and some neurons in motor cortex respond to visual and somatosensory cues (Ramkumar et al., 2016; Siniscalchi et al., 2016; Suminski et al., 2010; Fetz et al. 1980). While sensory and contextual signals might be useful for executing tasks, it remains unknown whether this non-motor-related M2 activity is routed to sensory regions via corollary discharge. If so, it raises important questions about the dynamic and reciprocal exchange of motor and sensory signals during behavior, and places important constraints on the mechanisms through which the brain learns and stores internal models that link action to outcome.

To address these questions, we taught mice to produce a simple forelimb movement that produced a simple sound while monitoring and manipulating neural activity in auditory and motor circuits in the cortex. We find that M2 encodes signals regarding movement as well as a movement’s acoustic consequences. Sound-related signals in M2 are routed back to the auditory cortex, suggesting a dynamic exchange of auditory- and motor-related signals during behavior. Finally, we find that sound-related signals in M2 are sensitive to expectation and experience. These experiments highlight the dynamic representation of sensory information in motor cortex and reveal that motor and sensory signals are bidirectionally exchanged during behavior.

## Results

### M2 neurons are active during a simple forelimb movement

We trained mice to make a silent, self-initiated lever movement, allowing us to first characterize how motor cortical cells encode an action that does not have an acoustic consequence (Fig. 1A). Mice began each self-initiated trial with the lever near their body (home position), were required to push the lever away from their body and past an uncued threshold (~15mm), and were rewarded with a drop of water when they returned the lever to the home position (Fig. 1B). Trials in which mice did not reach the movement goal and trials in which mice did not wait 500 milliseconds after finishing the prior movement were not rewarded. We trained mice on the lever behavior for 14 to 22 days and across training sessions mice increased the fraction of movements that resulted in a reward and pushed the lever more reliably toward the movement goal (Fig. 1C,D; Fig. S1A). By the end of training, mice made reliable lever movements that were clustered at reward threshold, lasted 275 milliseconds (median, +/−360 milliseconds, SD), and were rewarded on 40% (+/−2.6%, SEM) of trials that reached the movement goal (Fig. S1B,C).

**Figure 1.**
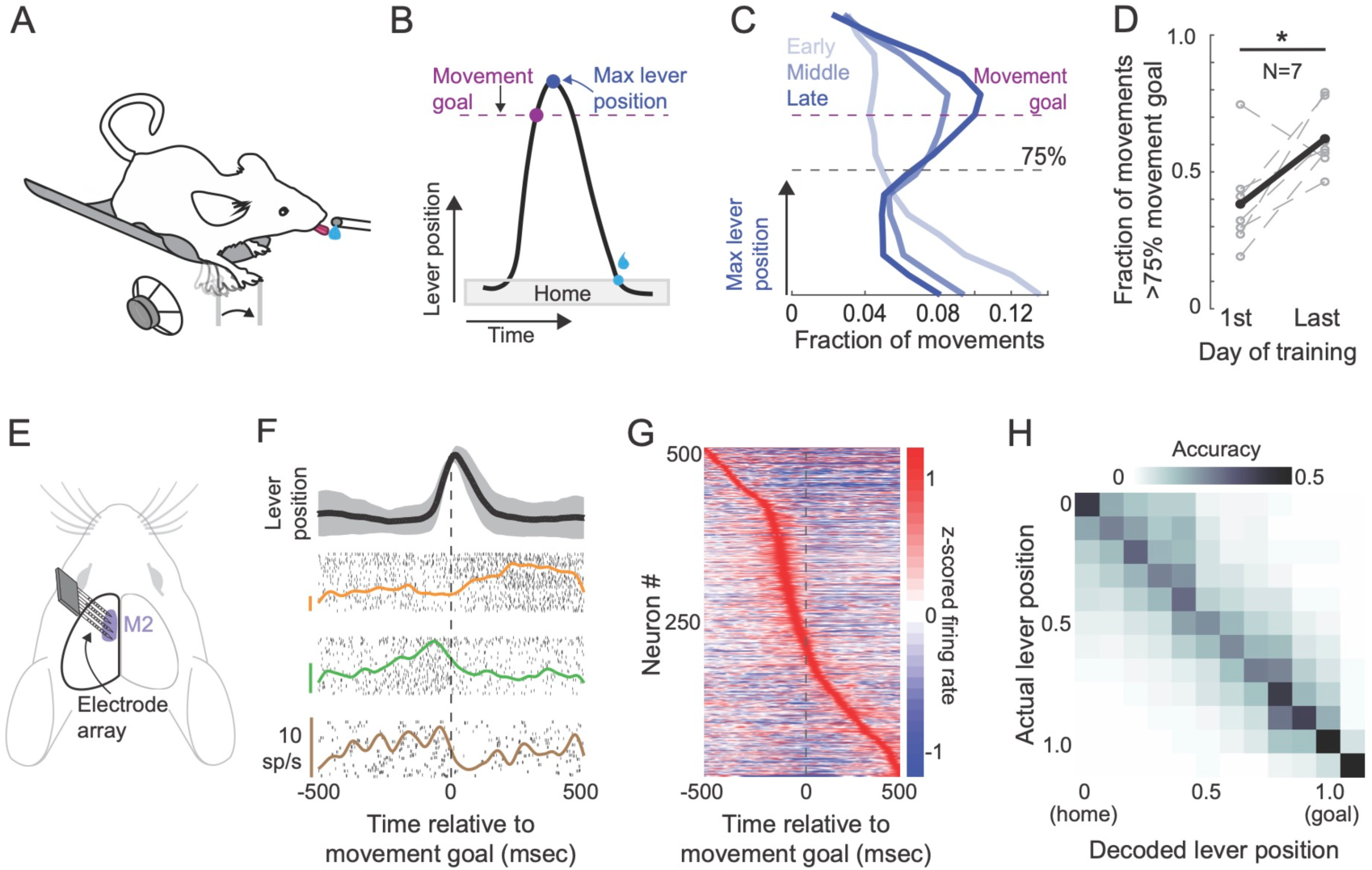
M2 neurons are active during a simple forelimb movement. (A) Schematic of a head-fixed mouse pushing a silent lever to receive a reward. (B) Schematic of a single trial, in which mice are required to push a lever with their right forelimb out of a home position past the movement goal, and then return the lever to the home position to receive a water reward. (C) Distribution of peak lever positions for an example mouse on three different days, sampled from early, middle, and late in learning. (D) Quantification of the fraction of lever presses that reached greater than 75% of the movement goal on the first versus last day of training across all mice. (E) Schematic of extracellular neural recordings from left M2 with a 128 channel electrode array. (F) Example lever trajectory (top) and raster plots with peri-stimulus time histograms for three example neurons, aligned to the movement goal. Shading over lever trajectory shows standard deviation. (G) Heatmap of z-scored activity of all recorded neurons (N = 7, n = 504). (H) Confusion matrix showing decoding accuracy from M2 neural activity. *p<0.05, **p < 0.005, ***p<.0005

Neurons in the secondary motor cortex (M2) are active during movements including forelimb movements and locomotion and are implicated in motor-sensory communication during sound-generating behaviors (Peters et al., 2014; Schneider et al., 2014; Schneider et al., 2018). Therefore, following behavioral training, we made acute electrophysiological recordings from M2 as mice operated the lever (Fig. 1E). Consistent with previous reports, we found that the majority of M2 neurons were active at some moment of the lever behavior, either in anticipation of, during, or after the lever movement (86%, 433 of 504 neurons) (Fig. 1F). As a population, M2 neural activity began to increase ~300 milliseconds prior to lever movement onset, peaked ~100 milliseconds prior to the movement goal, and gradually decayed throughout the remainder of the lever behavior (Fig. 1G). We trained a linear classifier using a subset of labeled neural data and measured how accurately we could decode lever position using withheld neural data (see Methods) (Fig. 1H). Decoding accuracy was significantly above chance throughout the outgoing phase of the lever press, consistent with M2’s role both in controlling movement and broadcasting reliable behavior-related information to other cortical structures. Altogether, these findings show that M2 activity reliably encodes a simple forelimb movement.

### M2 neurons respond following novel self-generated sounds

Many natural behaviors have acoustic consequences. Recent large-scale imaging and electrophysiology experiments show that signals related to movement and sensation are broadly distributed throughout the brain, including dorsal cortical areas involved in sensation, movement, and associative processing (Clancy, Orsolic, and Mrsic-Flogel 2019; Salkoff et al. 2020; Stringer et al. 2019; Musall et al. 2019). Using the same lever behavior, we next asked whether M2 activity changes when a previously learned silent movement begins producing a sound.

For 6 of the 7 mice trained on a silent lever, midway through electrophysiological recordings we added an experimentally controlled sound at a fixed phase of each lever movement (at the movement goal), and we recorded the activity of the same M2 neurons before and after the lever movement made a sound (Fig. 2A). The lever movements produced by the mouse in the 30 trials before introduction of the sound were indistinguishable from the lever movements after the introduction of the sound (Fig. 2B, Fig. S2A-C). Although the mouse’s lever movement did not change, many individual M2 neurons had a marked change in their activity immediately following the time of tone delivery (Fig. 2B). In the trials immediately following introduction of the self-generated sound, M2 activity continued to ramp up and peak early in movement and prior to the movement goal, but a prominent peak in population activity appeared during the 200 millisecond window following tone presentation (Fig. 2C). We compared neural activity before and after the introduction of the tone in two separate analysis windows, one just prior to and one just following the movement goal. We found that in the window of time just after the movement goal, which includes the time during which the sound was heard, there is a rapid and sustained increase in M2 activity that resembles a sensory-like response and persists across 30 or more trials (Fig. 2D). In contrast, in the window prior to the movement goal, which contains preparatory activity related to movement but not sound, M2 population activity remains unchanged after the introduction of the self-generated sound.

**Figure 2.**
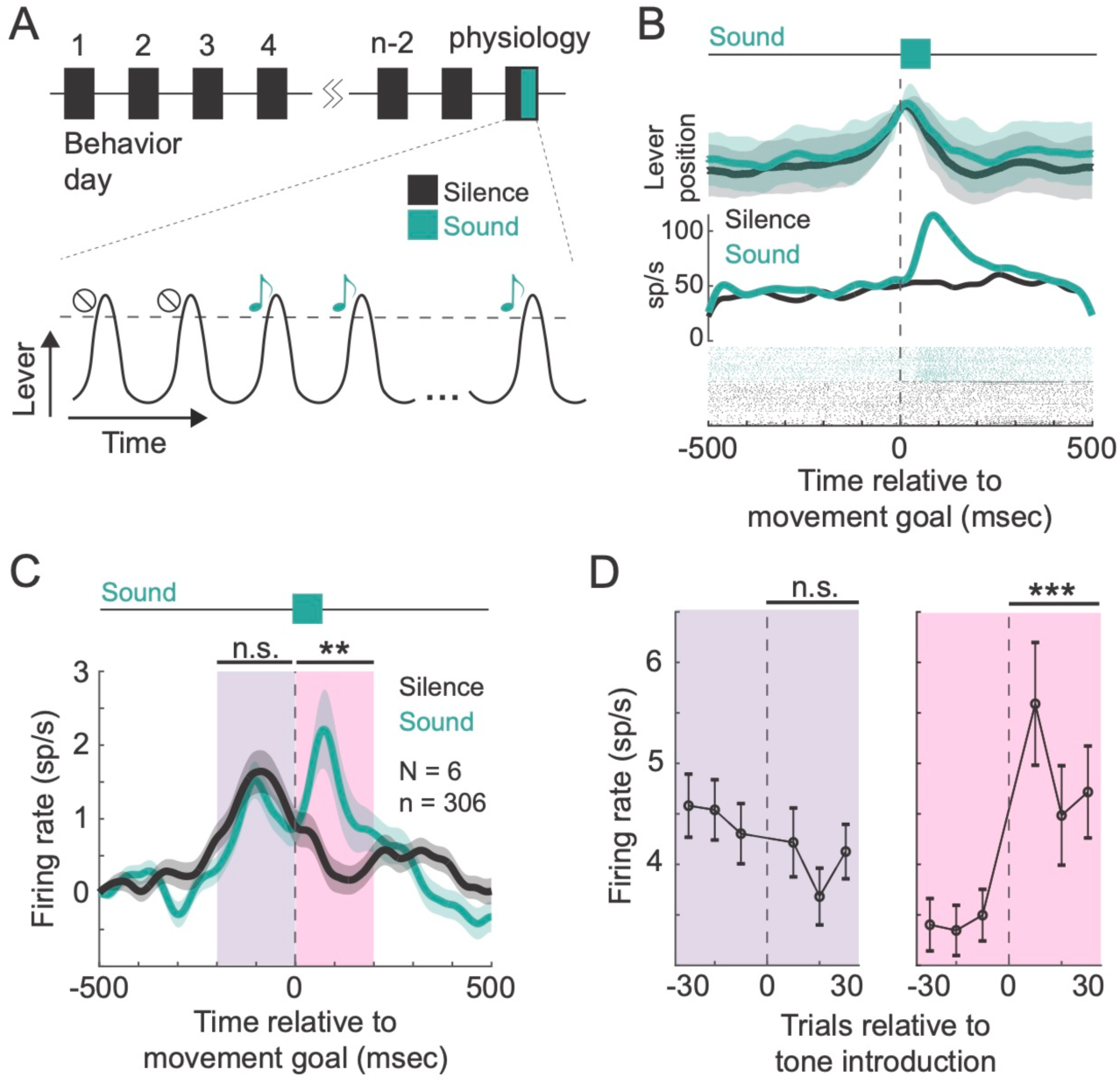
M2 neurons respond following novel self-generated sounds. (A) Schematic of the training and recording paradigm in silent-trained mice. The recording day consisted of an epoch of silent trials followed by an epoch of sound-generating trials. (B) Lever traces before and after introducing the self-generated tone (top), PSTHs of an example neuron (middle), and raster plots (bottom). (C) Population-averaged PSTH on the last 30 silent trials and the first 30 sound-generating trials, aligned to the movement goal. (D) Average firing rate in 30 trials before and after sound introduction (sliding 20-trial bins). Left shows activity in the 200 millisecond window before the movement goal was reached; right shows activity in the 200 millisecond window after the movement goal was reached. Shading shows standard deviation. *p<0.05, **p < 0.005, ***p<.0005

Across the recorded population, we found that 26% of M2 neurons (79 of 306) significantly changed their activity following the introduction of the tone. Of these 79 neurons, 33% were not responsive during silent movements, but only became responsive during sound-generating movements. Of the neurons that were significantly modulated by the introduction of the self-generated tone, the majority (58%) fired more action potentials during sound-generating compared to silent movements, while the other 42% had depressed responses during sound-producing movements. Taken together, these data suggest that in addition to reliably encoding movement kinematics, M2 neurons also encode the acoustic consequences of an action.

### M2 neurons are sound-responsive

Many M2 neurons are active following self-generated sounds, suggesting that M2 neurons integrate sound-related signals with ongoing motor-related activity. However, it remains unclear whether the peri-tone responses that we observe are sensory evoked or whether they reflect small behavioral responses to the sound. And if M2 responses are sensory-evoked, it remains unknown which regions of the brain provide sound-related input to M2. To address these questions, we removed the lever and presented tones of varying frequency (including the lever-associated frequency) during a passive playback epoch (Fig. S3A).

During passive sound playback, we found that many individual neurons had rapid and large responses following one or more tone frequencies (Fig. 3A). Across the population, a large fraction of M2 neurons (44%) were responsive to at least one passive sound and most neurons responded to multiple different frequencies (mode: 3 of 9; typically between 1 and 6) (Fig S3B).

**Figure 3.**
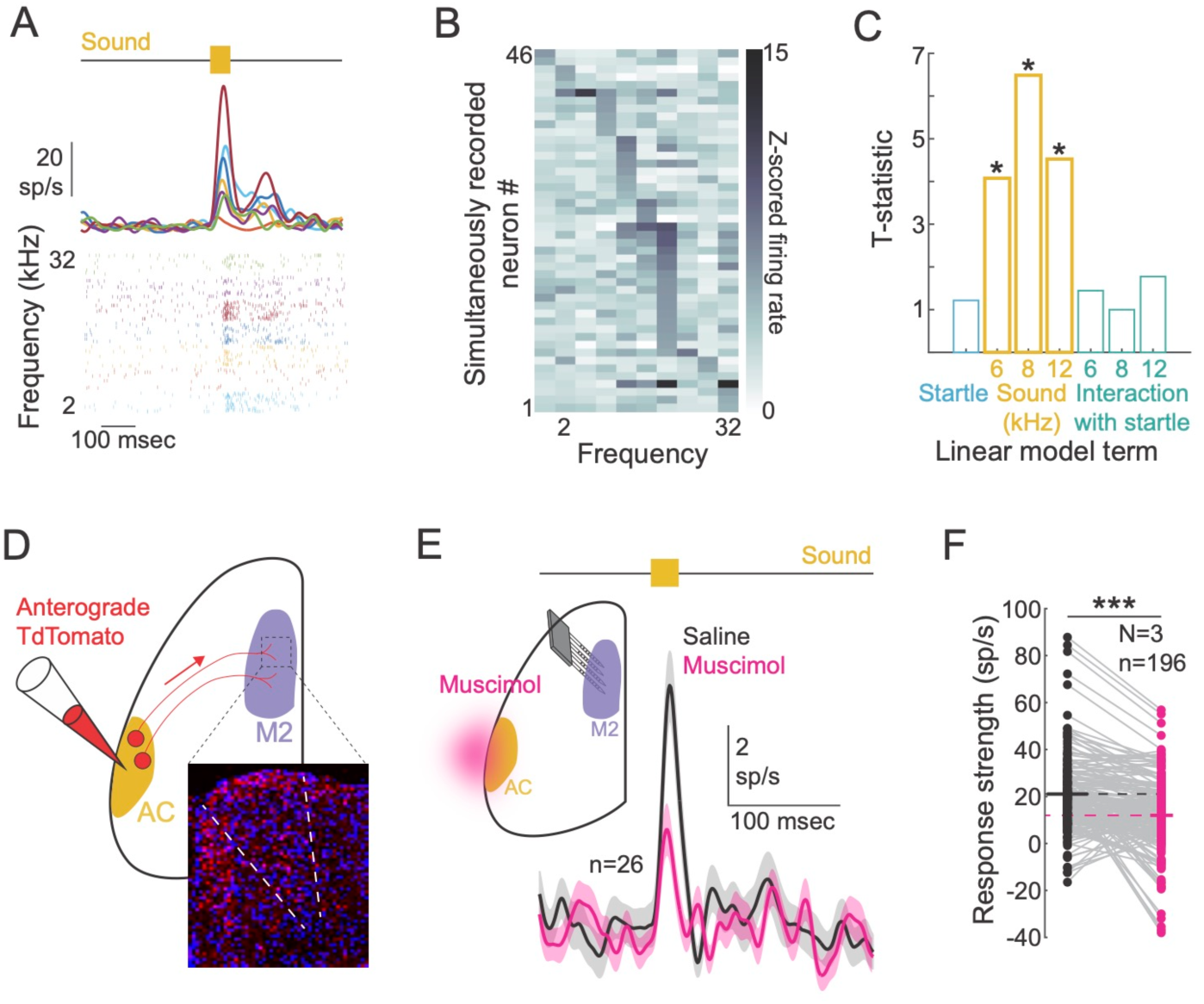
M2 neurons are sound-responsive. (A) Example neuron’s responses to passive sounds of varying frequency. (B) Heatmap of sound-evoked response magnitudes of simultaneously recorded neurons in an example mouse. (C) T-statistic results from a linear regression incorporating videography-captured startle movements, tone frequency, and interaction terms to predict trial-by-trial neural responses to passive sounds. (D) Schematic showing anterograde injection of AAV-TdTomato into auditory cortex and expression of TdTomato+ axon terminals in M2 (inset). (E) Schematic showing muscimol application to auditory cortex and simultaneous recordings in M2 during passive sound presentation. Average population activity of simultaneously recorded sound-responsive M2 neurons during saline and muscimol epochs from an example mouse. (F) Comparison of passive sound activity during saline versus muscimol trials in all recorded sound-responsive cells. *p<0.05, **p < 0.005, ***p<.0005

Several characteristics of sound-related responses in M2 suggest that they are sensory-evoked rather than being driven by reactionary movements. First, most sound-responsive M2 neurons had reliable responses at a short latency following sound onset (23% < 50 milliseconds; 71% < 100 milliseconds), consistent with responses observed in sensory cortex (Rummel et al., 2016; Guo et al., 2012) (Fig. S3C). Second, most sound-responding M2 neurons were responsive to only a subset of sound frequencies and simultaneously recorded neurons often had largely non-overlapping tuning curves (Fig. 3B). Finally, video analyses revealed that while some passively presented tones resulted in small startle movements, body movement alone is inadequate to account for the peri-tone activity that we observed in M2. As expected, some startle movements were accompanied by small increases in M2 activity, but the magnitude of startle movements in response to different frequencies was unrelated to the magnitude of neural responses to the same frequencies (Fig. S3D). Moreover, a linear model incorporating startle magnitude, tone frequency, and interaction terms revealed that responses to tones in M2 were better accounted for by sound frequency than by startle movements (Fig. 3C).

A possible route for acoustic information into M2 is via auditory cortex (AC), which in mice is reciprocally connected with motor cortical regions (Nakata et al., 2020, Nelson et al. 2013). We first verified that M2 receives direct input from the AC by injecting an anterograde tracer into AC and subsequently visualizing fluorophore-expressing terminals in M2, which were present across all layers, concentrated in superficial layers, and most prominent in ipsilateral M2 (Fig. 3D, Fig. S3E,F). Retrograde tracing from M2 confirmed the presence of M2-projecting cell bodies across layers in AC, concentrated in layer 2/3 (Fig. S3G-J). To determine whether these long-range projections from AC are involved in driving M2’s tone-evoked activity, we silenced AC using muscimol-soaked gelfoam applied directly to the surface of ipsilateral AC. Across the population, tone-evoked activity in M2 was decreased by 40% when ipsilateral AC was silenced compared to when AC activity was left intact (Fig. 3E). For many individual neurons (99 of 196 sound-responsive M2 neurons), tone-evoked responses were significantly weaker when AC was silenced (Fig. 3F). Together, these experiments reveal that many M2 neurons have sound-evoked responses that cannot be explained by movement and that M2 receives sound-related signals from auditory regions of the brain, including AC.

### M2 neurons have mixed selectivity for sound and movement

During a sound-generating behavior, population-level activity in M2 reflects a combination of movement-related activity and sound-evoked responses. We next investigated how these two types of signals – movement and sound – were encoded at the level of individual neurons. We measured whether each neuron was significantly activated during silent movements (movement-responsive), significantly responsive to passive sounds (sound-responsive), or both (Fig. S4A). Just under half of M2 neurons were sound responsive (44%), and of these cells, two thirds also had significant movement-related activity (Fig. 4A). We separated neurons into two groups, those that were responsive to the lever-associated frequency when presented passively (regardless of whether they were significantly movement responsive) and those that were responsive to movement but not to any sound. Analyzing the activity of these two populations separately, we found that sound-responsive neurons were more active during sound-generating behavior than were non-sound-responsive cells (Fig. 4B). This increased activity in sound-responsive cells was observed later in movement (after the sound was heard), early in movement (before the sound was heard), and during silent trials before the tone was introduced, suggesting that sound-responsive M2 neurons may be generally more active than M2 cells that are not sound responsive (Fig. 4C).

**Figure 4.**
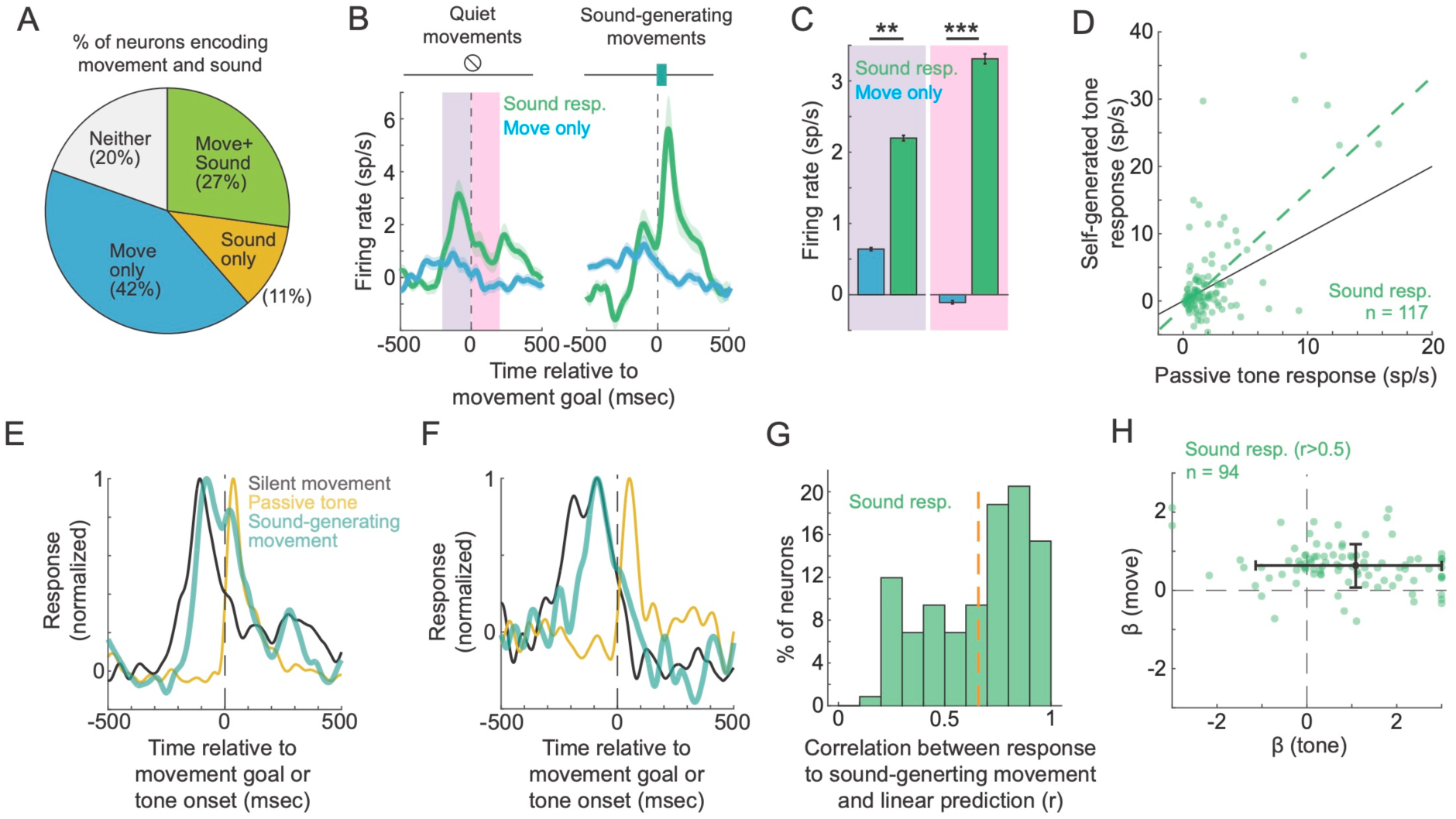
M2 neurons have mixed selectivity for sound and movement. (A) Pie chart showing proportion of M2 neurons that are responsive to movement, sound, both movement and sound, or neither. (B) Comparison of M2 activity in the movement-only and sound-responsive (sound only and move+sound combined) subpopulations during quiet (left) and sound-generating (right) movements. (C) Quantification of firing rates during quiet movements in the 200 milliseconds prior to when the lever reached the movement goal (purple) and the 200 milliseconds after the movement goal (pink), between movement-only (blue) and sound-responsive (green) cells. (D) Scatter plot of each neurons’ firing rates to self-generated tones and passive tones. (E) Activity of an example neuron whose activity during sound-generating movements (green) is well modeled as a linear sum of movement (black) and sound (yellow) responses. (F) Activity of an example neuron that responds to passive tones (yellow) but does not respond to self-generated tones (green) (G) Distribution of correlation coefficients between responses to sound-generating movements and the output of a linear model incorporating sound alone and movement alone, specifically for sound-responsive M2 neurons. (H) Scatter of tone and movement weights from the linear model (for neurons with r>0.5). *p<0.05, **p < 0.005, ***p<.0005

We next identified neurons that had a significant change in activity during sound-producing compared to quiet behaviors. Of these neurons that respond to self-generated sounds during movement, the majority also had a significant response when the lever sound was presented passively (54%). Across the sound-responsive M2 population, the magnitude of a neuron’s response to self-generated tones was correlated with its response to the same tone heard passively (r^2^ = 0.38, p < .001) (Fig 4D). These data further corroborate our observation that M2 sound responses are sensory-evoked rather than behavior-related.

Given the ubiquity of movement- and sound-related signals in M2, we next asked how these two signals interacted during sound-generating behaviors. For each neuron, we fit a linear model that compares the weighted sum of neural activity during quiet movements and passive sound playback to the activity measured during sound-generating movements. For many M2 neurons, activity during sound-generating movements could be well modeled as a linear sum of neural activity during silent movements and neural responses to passive tones (Fig. 4E,F). Focusing on the passive sound responsive population, the average correlation between neural responses to sound-generating movements and the linear model was 0.64 (Pearson correlation), including 70% of neurons with correlation coefficients greater than 0.5 (Fig. 4G). For these linear movement-sound neurons (r>0.5), the coefficients for the movement term of a linear model were generally positive in sign, consistent with neurons encoding movement during both quiet and sound-generating behaviors (Fig. 4H). The coefficient for the tone term was also positive on average, but was far more variable than the movement coefficient. While many M2 neurons could be well accounted for by a linear model, still other neurons were poorly fit by a linear model (30%), revealing a heterogeneity in how M2 neurons encode movement, sound, and the intersection thereof. Overall, the combination of movement and sound coefficients for linear M2 neurons were broadly distributed and reminiscent of the mixed selectivity for diverse task variables that has been observed in frontal cortex during cognitive tasks (Rigotti et al. 2013, Asaad and Miller, 1998; Hussar and Pasternak, 2009).

### Sound-evoked responses in M2 are relayed to auditory cortex

In addition to its role in behavior, movement-related activity in M2 is routed to sensory regions of the brain, including the auditory cortex, where these corollary discharge signals modulate sound-evoked activity (Nelson et al., 2013; Schneider et al., 2014). Corollary discharge signals are typically presumed to convey stable movement-related information to sensory areas regardless of a movement’s sensory outcome. However, given that M2 activity during sound-generating behavior reflects a combination of movement- and sound-related signals, we wondered whether M2 also relayed sound-related signals back to auditory cortex.

We injected an AAV2 expressing the optical neuronal actuator channelrhodopsin (ChR2) into M2, resulting in expression of ChR2 locally within M2 as well as in M2 axon terminals in the auditory cortex and other M2 targets (Fig. 5A). We then made electrophysiological recordings in M2 during silent and sound-generating behaviors and in response to passive sound playback. During these same recordings we optically activated ChR2-expressing axon terminals in the auditory cortex while monitoring antidromic action potentials in M2, allowing us to photo-identify M2 neurons that putatively send axons directly to auditory cortex, as well as many simultaneously recorded neurons that were not activated by terminal stimulation (Fig. 5B,C).

**Figure 5.**
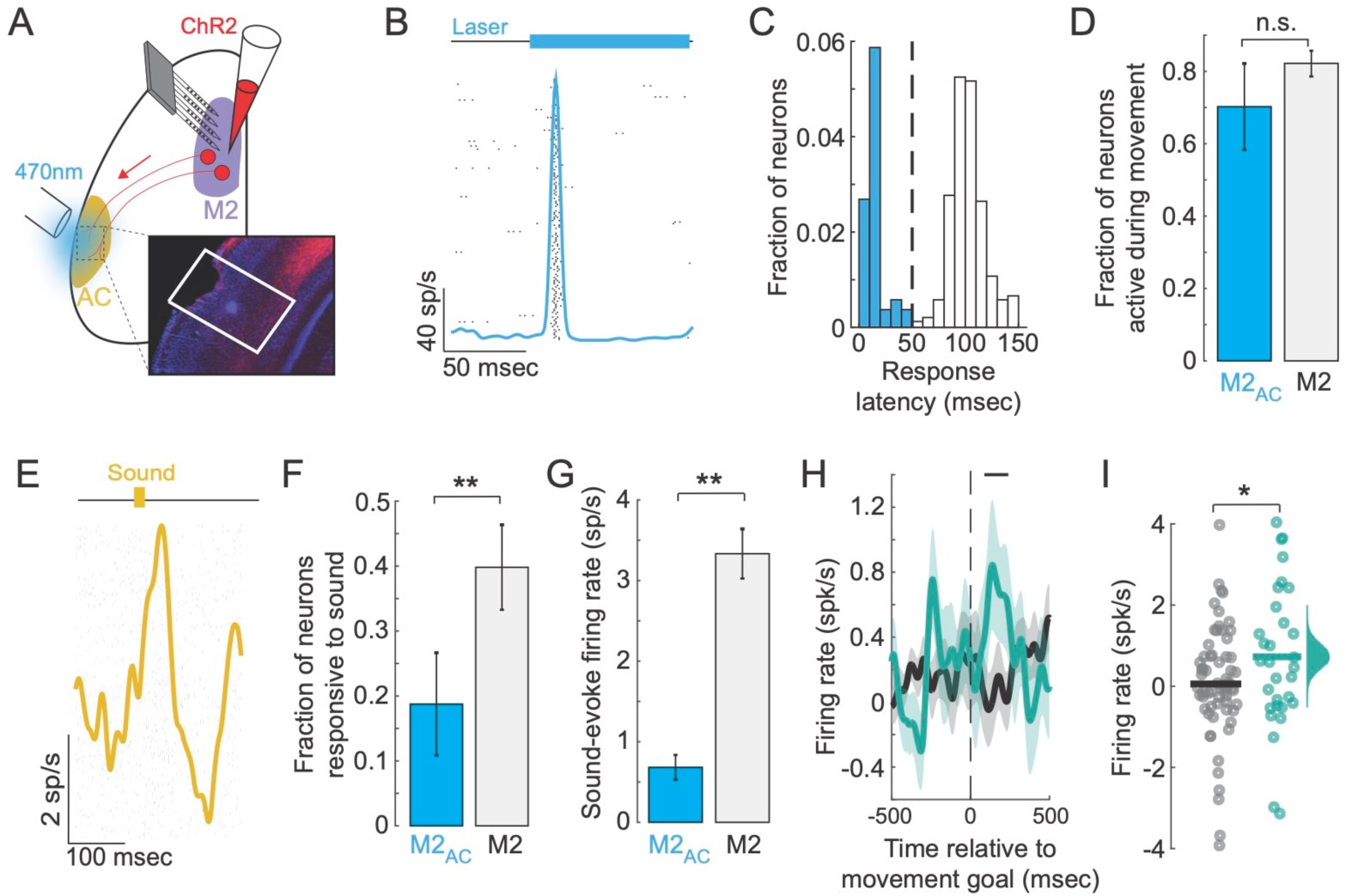
Sound-evoked responses in M2 are relayed to auditory cortex. (A) Schematic of optogenetic-tagging, where ChR2 is injected into M2 and an optical fiber is placed over the exposed auditory cortex. Inset: ChR2+ M2 terminals in AC with a small lesion where the optical fiber was positioned, made after the experiment. (B) Example photo-tagged AC-projecting M2 neuron (M2_AC_) with short latency antidromic activity following laser stimulation over auditory cortex. (C) Distribution of response latencies to laser stimulation, with blue indicating putative M2_AC_ neurons (latency <= 50ms) and open bars indicating non-tagged neurons (M2). (D) Fraction of movement-responsive neurons for M2_AC_ and M2 populations. (E) Passive sound-evoked response of an example M2_AC_ neuron. (F) Fraction of sound-responsive neurons for M2_AC_ and M2 populations. (G) Sound-evoked firing rate for M2_AC_ and M2 populations. (H) Average activity of M2_AC_ neurons on silent trials (black) versus trials that produced a tone (green). Black horizontal line indicates the window of analysis for panel I. (I) Firing rates of M2_AC_ neurons in a 200 millisecond window following the movement goal (and sound presentation on toned trials). Histogram shows bootstrapped estimates of the mean for sound-generating trials. *p<0.05, **p < 0.005, ***p<.0005

Like the general M2 population, AC-projecting M2 neurons contained multiple different classes of functionally identified neurons, including neurons that were responsive during silent movements (70%) (Fig. 5D) and neurons that were responsive to passively heard tones (19%) (Fig. 5D,E, Fig. S5A). The proportion of AC-projecting M2 neurons that were sound-responsive was significantly smaller than for the general M2 populations (40%) and the AC-projecting M2 neurons had weaker sound-evoked responses compared to the general M2 population (Fig. 5F,G), but these differences may reflect a smaller sample size for the AC-projecting M2 population (64 vs. 440 neurons).

During sound-generating behaviors, the AC-projecting M2 population had evoked activity that was aligned to sound onset, resulting in an increased firing rate during sound-generating movements compared to silent movements (Fig. 5H). This change in population-level activity was recapitulated at the level of individual neurons, where there was a significant increase in neural activity just after the tone was heard (Fig. 5I, Fig. S5B). These data reveal that, like the general M2 population, M2 neurons that project to the auditory cortex have movement-related activity and are responsive to passive sounds. Moreover, these AC-projecting M2 neurons send sound-related signals to the auditory cortex during sound-generating behaviors, showing that the corollary discharge signals routed from M2 to AC reflect a combination of movement and the sensory consequences of movement.

### M2 neurons do not respond to predictable self-generated sounds

Many behaviors produce the same sound every time they are executed, making their acoustic consequences highly predictable. Here, we have thus far only focused on M2 responses during movements that are silent or have recently acquired a new acoustic outcome. We therefore next asked whether M2 encoding of self-generated sounds changes when the sound becomes predictable based on its long-term association with a specific movement. To assess this, we trained mice on a sound-generating lever for 13 to 20 days and then made recordings in M2 (Fig. 6A, Fig. S6A). These sound-trained mice learned the behavioral task at a similar rate as silent-trained mice and had lever kinematics that were indistinguishable from those of silent-trained mice (Fig. 6B,C). The trial durations, fraction of movements that reached the movement goal, and fraction of goaled movements that received reward, were also similar to those of silent trained mice (Fig S6B-C).

**Figure 6.**
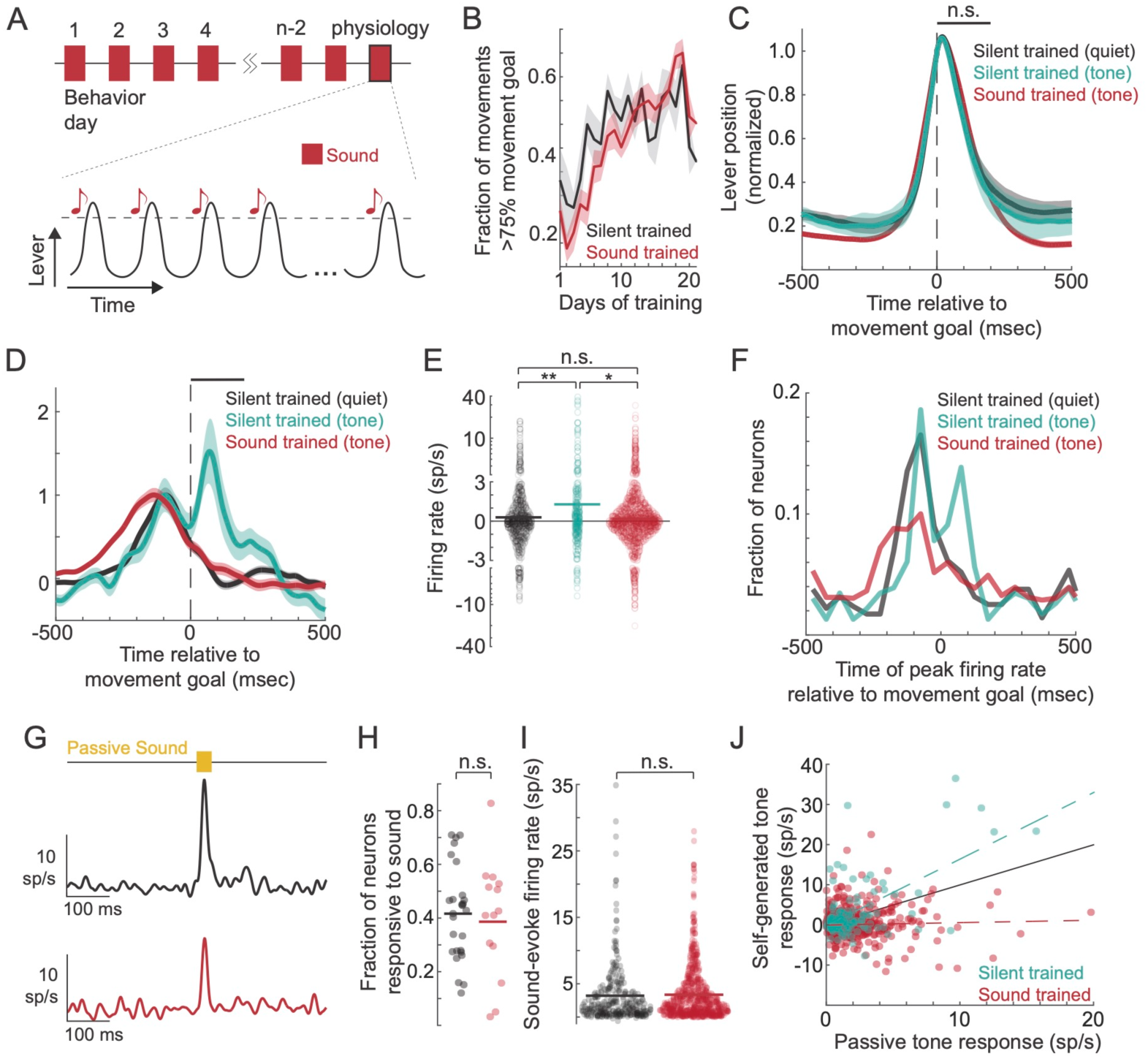
M2 does not respond to predictable self-generated sounds. (A) Schematic of the training and recording paradigm in tone-trained mice, with tones heard throughout training and on the day of electrophysiology. (B) Learning curves for silent-trained and tone-trained mice. (C) Average lever trajectory during physiology recordings for silent-trained mice operating a silent lever (black), silent-trained mice operating a sound-generating lever (green), and tone-trained mice operating a sound-generating level (red). (D) Comparison of M2 population activity centered on the movement goal across the three conditions. Black line indicates the window of analysis in 6E. (E) Comparison of cells’ firing rates across the three conditions. (F) Distribution showing the time of peak firing rates for each neuron across the three conditions. (G) Example sound-responsive neurons from a silent-trained mouse (black) and tone-trained mouse (red). (H) Fraction of tone responsive neurons across recordings in silent-trained and tone-trained mice. (I) Magnitude of the tone-evoked activity in the sound-responsive populations in silent-trained and tone-trained mice. (J) Scatter plot of each neurons’ firing rates to self-generated tones and passive tones, for both silent-trained (green) and tone-trained (red) mice. *p<0.05, **p < 0.005, ***p<.0005

Following behavioral training, we recorded from M2 neurons as mice made lever movements that produced a predictable tone. Similar to mice trained in silence, we could reliably decode the lever position from M2 activity in tone-trained mice (Fig. S6D). We also found a similar number of M2 neurons active during the lever movement in tone-trained mice compared to silent-trained mice (84%, 1357 of 1609 neurons) and similar proportions of cells responsive to movement, sound, or both (Fig. S6E).

However, unlike in mice who heard a self-generated sound for the first time during electrophysiology, M2 neurons in tone-trained mice showed no noticeable sound-evoked activity during behavior (Fig. 6D). On average, M2 neurons in tone-trained mice had equivalent firing rates as mice trained on and operating a silent lever, and significantly weaker firing rates than mice hearing a tone for the first time (Fig. 6E). Across the population, the time of peak activity during behavior for M2 neurons was biased before the movement goal (and tone time) in mice making silent movements and in mice hearing a predictable sound. In contrast, for mice hearing an unexpected self-generated sound, a substantial fraction of neurons had their peak activity after the time of the tone (Fig. 6F). As observed in silent-trained mice, we found that tone-responsive cells were significantly more active during behavior than were non-sound-responsive cells, even prior to the tone being heard (Fig. S6F).

Given the lack of responses to self-generated sounds in tone-trained mice, we next tested whether the M2 neurons we recorded from in tone-trained mice were capable of responding to tones in the passive condition. As we had seen in silent-trained mice, individual M2 neurons in tone-trained mice had strong sound-evoked responses with similar sound latencies (Fig. 6G; Fig. S6G). Across experiments, we found a similar number of passive sound-responsive neurons in M2 of silent-trained and tone-trained mice, and tone-responsive neurons had similar tone-evoked firing rates in response to passive sounds (Fig. 6H,I). However, unlike in mice hearing a self-generated sound for the first time, in tone-trained mice we observed no correlation between neurons responses to self-generated sounds and the same sounds heard passively (Fig. 6J). Therefore M2 neurons in sound-trained mice are capable of responding to sounds, but during movements with a predictable acoustic consequence, M2 responses appear to be gated off.

Finally, we injected ChR2 into M2 of tone-trained mice, allowing us to photo-identify AC-projecting M2 cells during electrophysiology. We found that a significantly larger fraction of AC-projecting cells were active during the behavior in tone-trained mice compared to silent-trained mice (87% vs 69%, z-test = −3.35, Fig. S6H). Following the self-generated tone, the AC-projecting M2 neurons in tone-trained mice had a prominent negative dip in their activity, in contrast to the same population’s positive-going response following self-generated sounds in silent-trained mice (Fig. S6I). This dip in activity immediately following the tone in tone-trained mice appeared to be driven by the tone, as it was absent on interleaved trials that did not produce a sound because they nearly reached, but did not reach, the movement goal (Fig. S6J). Interestingly, in both silent-trained and tone-trained mice, AC-projecting M2 neurons exhibit an increased firing rate when mice hear something unexpected during behavior: in silent-trained mice, responses are larger when movements unexpectedly produce a tone, and in tone-trained mice, responses are larger when mice unexpectedly hear silence (Fig. S6I,J).

### Population dynamics in M2 reflect movement, sensation, and experience

Population dynamics in motor and sensory cortex tend to be constrained along distinct manifolds and the dimensions that encode motor and sensory signals tend to be largely orthogonal to one another (Kaufman et al., 2014, Elsayed et al. 2016, Stringer et al., 2019). To quantify population geometry in M2 during sound-generating behaviors, we performed a singular value decomposition (SVD) and projected trial-averaged neural population data into a two-dimensional space that accounted for 59% of the variance (Fig. 7A). Using this low-dimensional representation of M2 activity, we aimed to determine how movement- and sound-encoding dimensions interact at the population level during behavior, and whether movement- and sound-encoding dimensions are influenced by expectation.

**Figure 7.**
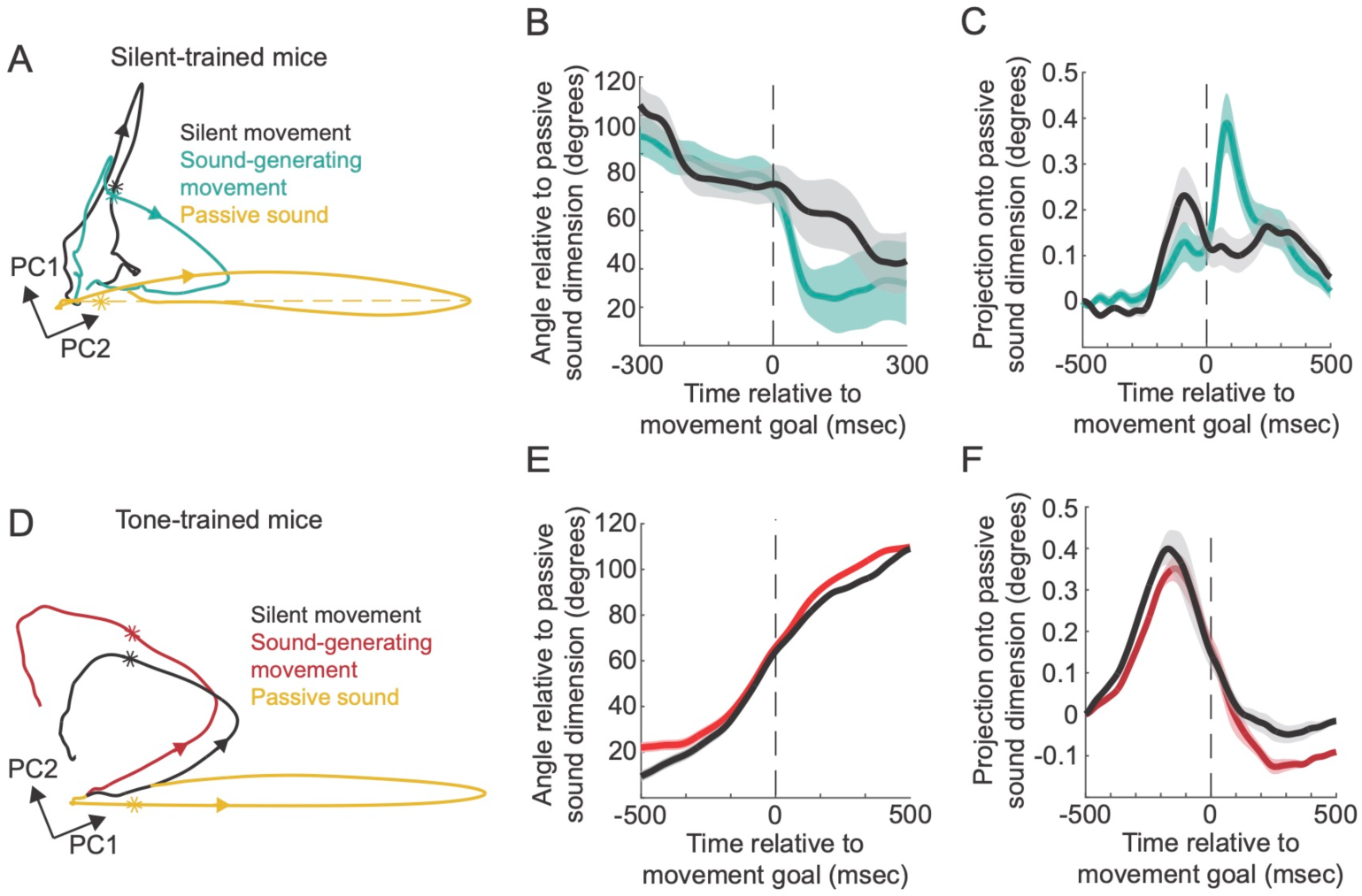
Population dynamics in M2 reflect movement, sensation, and experience. (A) Neural trajectories of silent-trained mice in a two-dimensional PCA subspace, comparing activity of the M2 population on silent trials (black), during passive sound playback (yellow), and during sound-generating movements (green). Stars indicate the time of the movement goal or tone. Arrows indicate the direction of the trajectory forward in time. Yellow dashed line shows the vector pointing in the sound-encoding dimension. (B) Quantification of the instantaneous angle between the trajectory for sound-generating movements and the vector pointing in the sound-encoding dimension (yellow dashed line in panel A). (C) Projection of the trajectory for sound-generating movements onto the vector pointing in the sound-encoding dimension. (D) As in panel A, but for tone-trained mice. Red shows activity during sound-generating movements. (E and F) As in panels B and C, but for tone-trained mice.

We first compared neural activity during silent movements and during passive sound playback. Comparing the temporal trajectories of these two trial types revealed that the population activity during passively played sounds was largely orthogonal to population activity during silent movements (65 degrees). We next quantified the temporal trajectory of population activity in silent-trained mice hearing a new self-generated sound. To quantify the degree to which self-generated sounds impact population dynamics in M2, we projected the M2 population activity onto a vector that reflects the sound-encoding dimension (Fig 7A, dotted yellow line) and we also compared the angle of the instantaneous population vector with the vector along the sound-encoding dimension (Fig. 7B,C). There is a weak interaction between the movement and sound subspaces early in behavior (before the sound is heard), manifesting as an interaction angle less than 90 degrees and a non-zero projection of movement onto the sound-encoding dimension. The M2 trajectory begins interacting more prominently with the sound subspace at the time of the lever-produced tone, resulting in a larger projection onto the sound-encoding dimension and a decreased angle relative to this dimension (Fig 7B,C). These population geometry analyses are consistent with our single-cell and population PSTH-based analyses, showing that M2 activity is influenced by both movement and sensory experience.

We next analyzed the temporal trajectories for M2 populations in tone-trained mice (Fig. 7D). In contrast to silent-trained mice, the dimensions encoding movement and sound were less separated from one another, and even during silent movements there was a substantial projection onto the sound-encoding dimension (Fig. 7E,F). In tone-trained mice, M2 population dynamics were nearly identical during sound-generating and silent movements, exhibiting a maximum projection onto the sound dimension early in movement (~200 msec before the sound occurred) and showing no increased alignment with the sound dimension after the sound was heard (Fig. 7E,F).

These results suggest that with experience, the sound- and movement-encoding dimensions in M2 become less independent; that the presentation of sound has a qualitative impact on ongoing population dynamics in silent-trained but not tone-trained mice; and that following extensive experience, movement signals interact with sound-encoding dimensions early in movement, consistent with expectation-like processing in M2.

## Discussion

We designed a behavioral paradigm in which the acoustic consequence of a simple forelimb movement was under experimental control. Using this behavior, we found that neurons in M2 had activity patterns that reflected behavior and its sensory consequences. M2 neurons are sound responsive outside of behavior, in part due to input from the auditory cortex, and sound-related signals in M2 are routed back to the auditory cortex during sound-generating behaviors. After extensive experience with a sound-generating behavior, M2 neurons no longer respond to self-generated sounds yet retain their responses to passive sounds. These changes in single-cell responses can be observed at the population level, where experience alters the geometry of cortical coding during sound-generating behaviors, leading to a realignment of sensory- and movement-encoding dimensions. Together, these findings reveal the embedding of sensory and expectation signals in motor cortical activity.

We find that auditory-related activity is prominent in M2 and is sensitive to experience and expectation, consistent with M2’s extensive connectivity with other brain areas. M2 receives information from sensory, parietal, orbital and retrosplenial cortices, as well as thalamus, and relays motor information to the spinal cord, superior colliculus and subcortical nuclei (Barthas and Kwan, 2015). Our data reveal rapid and strong sound-evoked responses in M2 that arise in part from the auditory cortex and cannot be accounted for by small reactionary movements. Instead, we find that M2 responses to self-generated sounds are correlated with the same neurons’ responses to passive sounds and that as a population, M2 neurons exhibit a heterogeneous combination of movement- and sound-related signals, consistent with the mixed selectivity that has been previously observed in frontal cortex (Rigotti et al., 2013).

While the current study was not designed to identify the function of auditory signals in M2, we note that such signals may be important for auditory-guided motor learning. For example, when the pitch of a subject’s voice is artificially augmented using closed-loop headphones, humans, marmosets, and songbirds adjust their pitch so what they hear through the headphones matches what they expect to hear (Sober & Brainard, 2009; Elman JL, 1981; Eliades & Tsunada, 2018). In marmosets, these changes in behavior are correlated with auditory cortical activity, suggesting that auditory cortex may route sound-related signals to vocal control areas during behavior (Eliades & Tsunada 2018). M2 also shares homology with the premotor nucleus HVC in songbirds, which is both essential for song production and receives input from auditory centers of the telencephalon (Prather et al., 2008). Normal song learning requires intact auditory feedback as well as auditory input to singing-related structures of the songbird brain (Mandelblat-Cerf et al., 2014; Konishi, 2004). Whether auditory signals in mouse M2 also facilitate auditory-guided learning will require new behavioral paradigms in which mice adjust their behavior based on acoustic feedback.

We find that sensory-evoked signals in M2 are routed back to the auditory cortex, Models of predictive processing and motor-sensory learning posit that corollary discharge signals are passed through an internal model, resulting in an efference copy, which anticipates the specific sensory consequences of an action (Keller & Mrsic-Flogel, 2018). While the cellular and circuit locus of internal models remains largely unresolved, our data support a shift away from a static, movement-centric view of corollary discharge, toward a more dynamic and multi-modal view. Our findings are consistent with previous work that has observed that top-down input to sensory areas can be impacted by sensation, including mixed visual and motor information in corollary discharge signals relayed by superior colliculus during saccades (Sommer and Wurtz, 2004; Subramanian et al, 2019) and experience-dependent changes of top-down, behavior-related inputs to visual cortex during locomotion (Leinweber et al., 2017).

Following extensive experience with a sound-generating behavior, we find that M2 neurons cease responding to self-generated sounds. Suppression of neural responses to self-generated sounds is a prominent feature of auditory cortex, and M2 receives some sound-related input from auditory cortex. Interestingly, rather than being abolished, sound-evoked responses in A1 during the same lever behavior are suppressed by ~50% following extensive training (Audette et al., 2022). These data suggest that the loss of input from the auditory cortex alone is likely inadequate to explain the lack of sound-responsiveness in M2 of experienced mice. Instead, it is possible that sound-evoked responses are gated off locally within M2, reminiscent of auditory responses in songbird HVC. Under anesthesia, HVC neurons are responsive to playback of a bird’s own song. These responses largely go away when the bird is awake, even though Field L, an auditory cortical area that provides input to HVC, remains responsive (Schmidt and Konishi 1998). Future studies are necessary to determine the mechanisms by which sound-evoked responses are gated off in M2 following extensive training.

M2 and neighboring anterior cingulate cortex have been implicated in predictive processing during behavior. We find that in silent trained mice, M2 neurons have larger responses when a movement makes a sound (and violates the mouse’s expectation) compared to when the same movement is silent (i.e. matches expectation). Conversely, in tone-trained mice, neural responses in M2 are weak when the movement produces a tone (matches expectation) and are larger on trials when a movement does not produce a tone (a violation). Expectation-like signals are also present in our analyses of population dynamics, where we observe an alignment of sound-encoding and movement-encoding dimensions following extensive experience with a sound-generating movement. Predictive processing during behavior has been extensively studied in sensory cortex (Schneider et al., 2018; Audette et al., 2022; Audette et al., 2023; Keller et al., 2012) and our data suggest that motor cortex may also be involved in forming expectations and encoding violations.

Taken together, these results build a foundation for further investigations into the loci of internal model storage in predictive processing, the function of sensory information in motor cortical circuits, and the nature of dynamic, multi-modal corollary discharge signals during behavior.

**Supplementary Figure 1.**
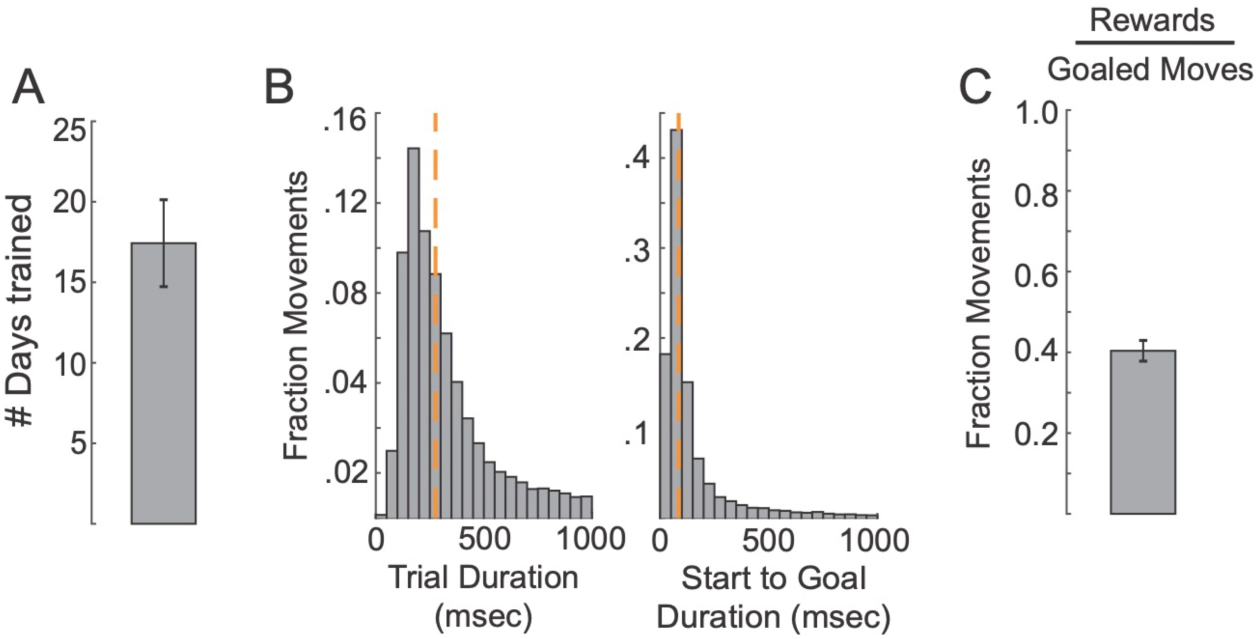
(A) Average number of days that mice were trained to push a silent lever. (B) Distribution of trial durations (left) and distribution of times from start (leaving the home position) to reaching the movement goal (right). Orange line indicates the median of the distribution. (C) Average percentage of movements that reached the goal threshold that were rewarded.

**Supplementary Figure 2.**
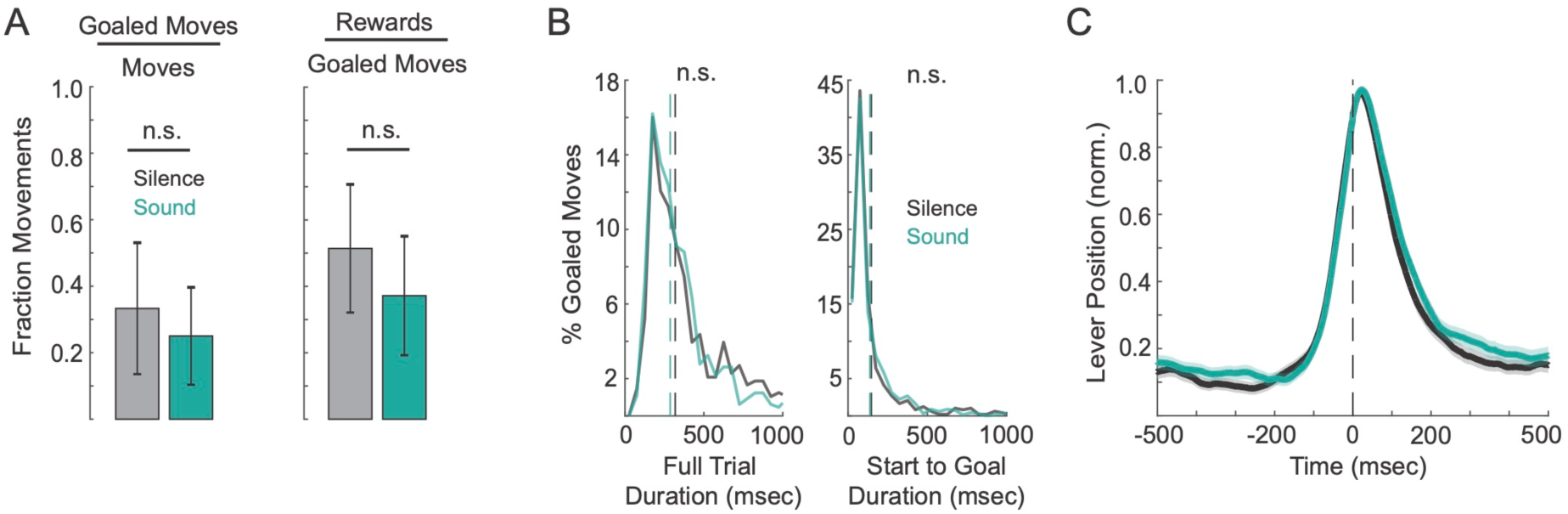
(A) Left: Fraction of movements that reached the movement goal on silent trials and sound-generating trials. Right: Fraction of goaled movements that were rewarded on silent trials and sound-generating trials. (B) Left: Distributions of movement durations for silent trials and sound-generating trials. Right: Distributions of the duration from leaving the home position to reaching the movement goal. (C) Grand average lever trajectories across mice on silent trials and sound-generating trials (N = 6).

**Supplementary Figure 3.**
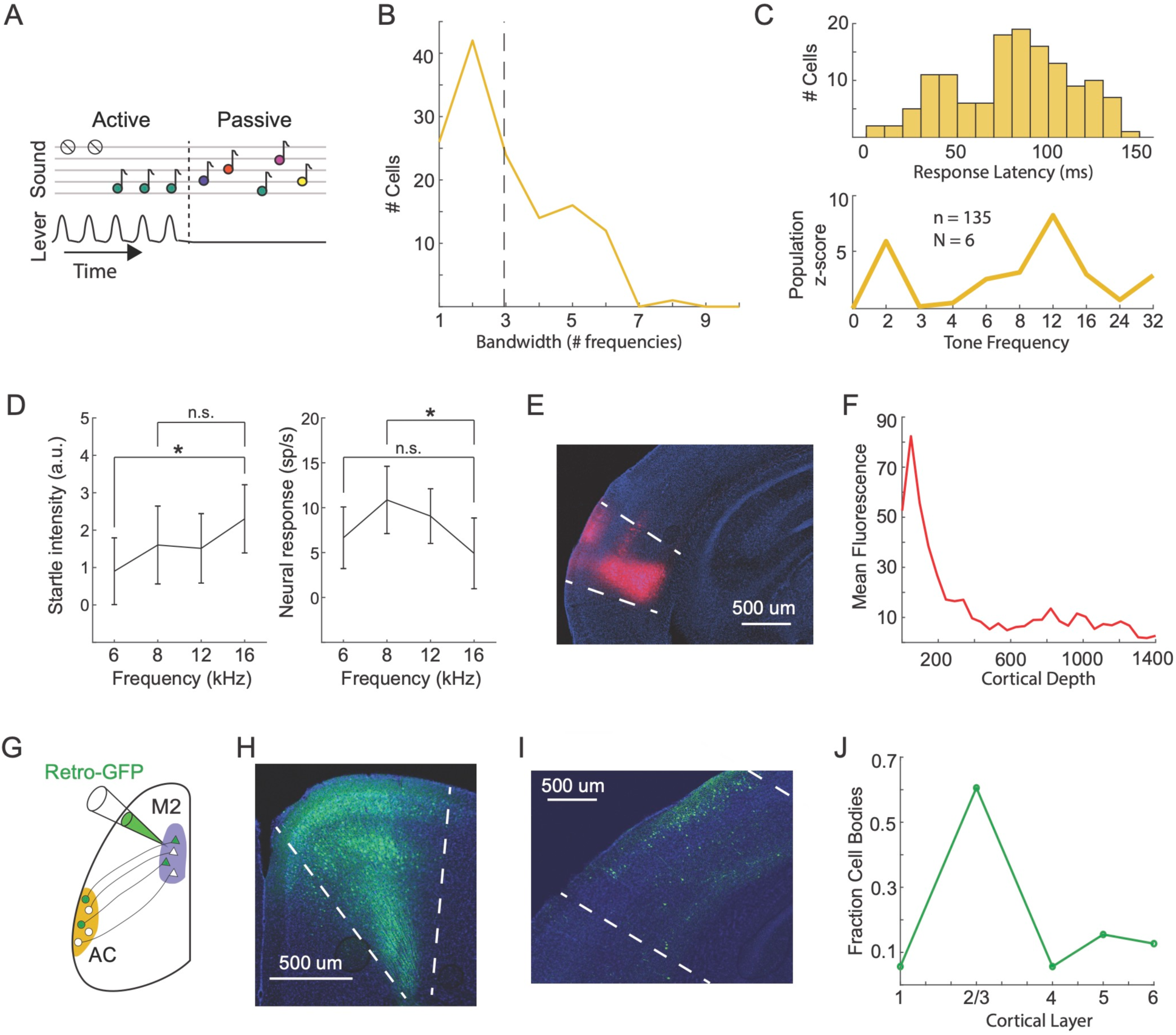
(A) Schematic of sounds presented passively to the mouse with the lever removed. (B) Bandwidth of individual neurons. (C) Distribution of response latencies to passive sounds (top) and population-averaged responses to all sound frequencies used (bottom). (D) Startle intensity of frequencies with the largest population-averaged responses (left) and population-averaged neural responses to the same frequencies (right), showing that frequencies with the largest startle response are not the same as those with the largest neural response. (E) Injection site of anterograde tracer into auditory cortex. (F) Quantification of terminal density in M2 following injection of anterograde tracer into auditory cortex. (G) Schematic showing injection of retrograde AAV-GFP into M2. (H) Injection site of retrograde tracer into M2. (I) Retrogradely labeled cell bodies in auditory cortex that project to M2. (J) Distribution of M2-projecting auditory cortex cell bodies, by cortical layer.

**Supplementary Figure 4.**
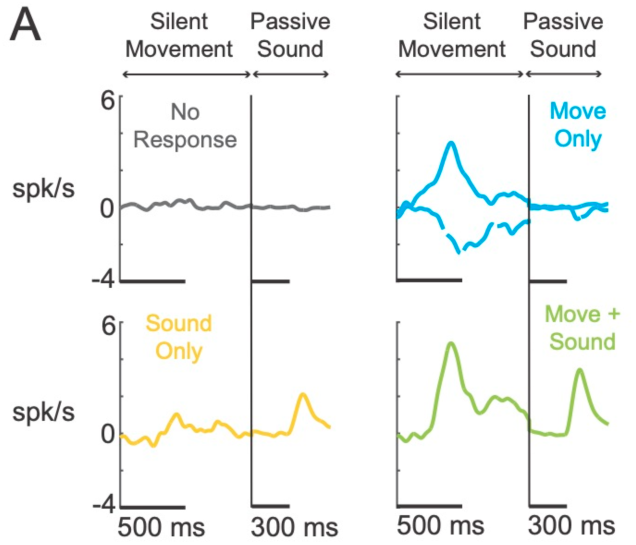
(A) Subpopulation-averaged neural responses during silent movements and in response to passive sounds, for each of 4 groups of M2 neurons described in Fig. 4A. For each group, the left PSTH shows responses to silent movements and the right PSTH shows responses to passive sounds. Neurons in the “Move only” category were separated based on whether they were positively (solid) or negatively (dashed) modulated during movement.

**Supplementary Figure 5.**
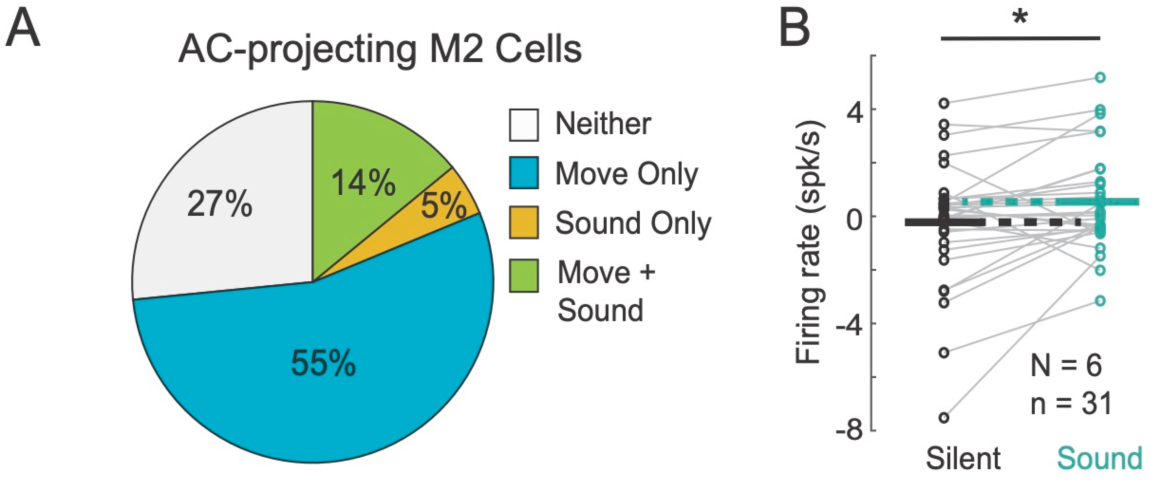
(A) Percentage of AC-projecting M2 neurons in silent-trained mice that are responsive to movement, sound, movement and sound, or neither. (B) Firing rate of simultaneously recorded AC-projecting M2 neurons on silent trials and sound-generating trials.

**Supplementary Figure 6.**
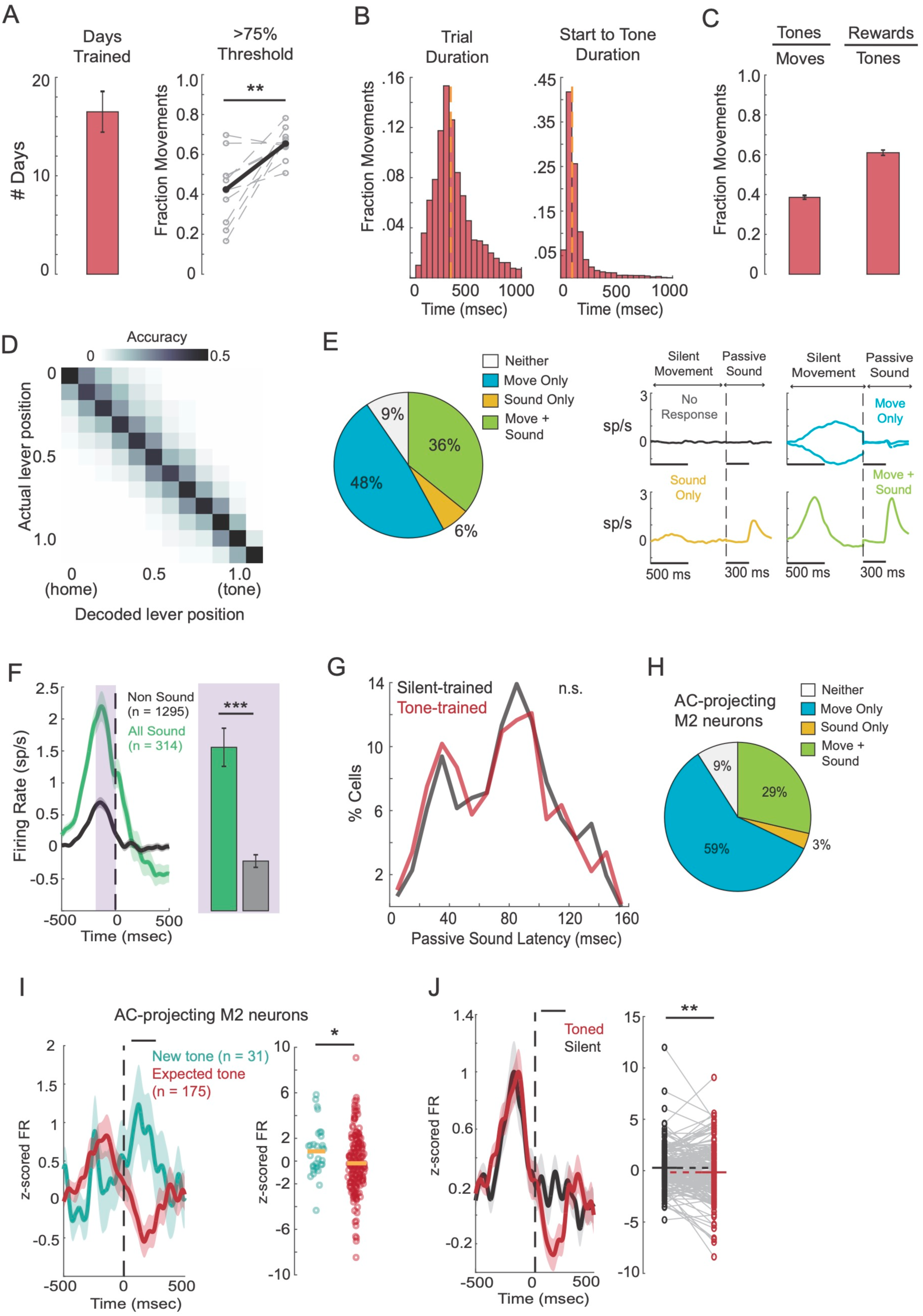
(A) Left: Average number of training days for tone-trained mice. Right: Fraction of movements that exceeded 75% the distance to the movement goal on the first and last day of training (N = 9 mice). (B) Left: Distribution of durations from leaving the home position to returning to the home position. Right: Distribution of time from leaving the home position to reaching the movement goal. Orange dashed lines show median. (C) Left: Proportion of movements that were sound-generating. Right: Proportion of sound-generating movements that received a reward. (D) Confusion matrix comparing actual lever position with lever position decoded from M2 activity in tone-trained mice. (E) Left: Percentage of M2 neurons in tone-trained mice that were modulated by movement, sound, both, or neither. Right: Subpopulation-averaged activity for neurons belonging to each category. (F) Left: Population PSTHs for sound-responsive (green) and movement-responsive (but not sound-responsive) neurons (black). Right: Average firing rates in the 200 millisecond window before the movement goal. (G) Distribution of latencies of tone-evoked responses in silent-trained and tone-trained mice. (H) Percentages of AC-projecting M2 neurons that respond to movement, sound, both, or neither. (I) Left: Population PSTHs for AC-projecting M2 neurons during sound-generating movements in silent-trained mice (green) and tone-trained mice (red). Right: Firing rates for the individual neurons that make up the populations on the left. (J) Left: Population PSTHs for AC-projecting M2 neurons recorded in tone-trained mice, on sound-generating trials (red) and quiet silent trials (black). Right: Firing rates of AC-projecting M2 neurons during silent and sound-generating movements in the window immediately after the sound was heard.

## Methods

### Animals

All experimental protocols were approved by New York University’s Animal Use and Welfare Committee. Male and female wild-type (C57BL/6) mice were purchased from Jackson Laboratories and were subsequently housed and bred in an onsite vivarium. All experimental mice were 2-4 months old and were kept on a reverse day/night cycle (12h day, 12 h night).

### Viral Injections and Electrophysiology Surgeries

For all surgical procedures, mice were anesthetized under isoflurane (3-4% in O2) and head-fixed in a stereotaxic holder (Kopf). Hair and skin were removed from the top of the head. For viral injections of channelrhodopsin, virus (AAV-CaMKIIa-hChR2-mCherry, UNC, titer 3.9e12) was injected into M2 (1.0-1.5mm AP, 0.5-0.7mm ML from bregma) at three depths (200um, 400um, and 600um), 100nL at each depth for a total injection volume of 300nL. A metal headplate (H.E. Parmer) was attached to the skull using a transparent adhesive (Metabond). Mice were treated with an analgesic (Meloxicam SR) and allowed to recover for 5 days prior to training. Post-training and 24 hours prior to electrophysiology, a small craniotomy was made to expose M2 at and around the injection site. For all optotagging experiments, a small craniotomy was also made over auditory cortex (~2mm diameter, −2.5mm AP, and ~4.2mm ML), which had been previously marked during the initial surgery. A final craniotomy was made above right visual cortex, where a silver-chloride ground electrode was positioned under the skull, atop the surface of the brain, and covered with Metabond. Exposed craniotomies were covered with a silicone elastomer (Kwik-Sil) and the mouse was allowed to recover in its home cage.

### Behavior Training and Analysis

We used a custom lever push behavior that produced closed-loop sounds, as previously described (Audette et al., 2022). All training and experiments were performed in a dark, sound-attenuating booth (Gretch-Ken) to minimize background sound and monitored in real-time via IR video. Water-restricted mice were head-fixed to the behavioral apparatus and presented with the lever and lickport after ~10 minutes of quiet acclimation. Mice were then allowed to self-initiate movements. For mice trained in silence, the movement goal was set ~12mm from home position. We also required the lever to remain in the home position (~+/− 3mm from rest) for >500 milliseconds prior to initiation for a movement to be considered valid. Valid movements that crossed the movement goal elicited a small water reward (5-10uL) when the lever returned to home position. For mice trained on a sound-generating lever, trial requirements remained the same as mice trained in silence, but auditory feedback was delivered for all movements that crossed the movement goal. If the lever crossed the movement-goal threshold multiple times within a single trial, the tone was delivered only upon the first crossing. Mice were trained on a single sound frequency, but the trained frequency varied across mice and ranged between 2 and 32 kHz. Auditory feedback was provided on all trials in which the lever crossed the movement-goal threshold, regardless of whether mice obeyed the inter-trial wait time requirement and received a reward.

Mice were trained one session per day for between 13 and 22 days prior to neural recording. Prior to the first training session, mice were water-restricted for 24 hours. In the early stages of training the movement-goal was nearer the mouse, and was incremented farther away during the first few sessions. The home position requirement (500 milliseconds) was omitted to allow the mice to quickly gain an understanding of the task. Within 3-5 sessions the movement goal was shifted to its final position and the inter-trial interval requirement for reward was implemented and gradually increased to 500 milliseconds, and these parameters were fixed for the remainder of training. Lever trajectories were analyzed post-hoc and across training sessions to assess mouse behavioral patterns and learning. To assess improvement on the task across sessions, we quantified the percentage of trials where the lever was displaced >75% of the reach threshold.

To quantify startle movements during sound playback, we extracted motion energy from video frames just prior to and just after sound playback (+/− 300 milliseconds). To calculate motion energy during each tone presentation, we first identified the video frame during which a tone was presented. Next, we cropped each frame to include only the mouse’s face and body, performed a pixel-by-pixel subtraction of frame N+1 from frame N (for each pair of subsequent frames), took the absolute value of each pixel difference, and summed across all pixels. We performed this computation separately for frames before sound playback and frames after sound playback. We then subtracted motion energy before the sound from motion energy after the sound, resulting in a change in motion energy relative to a pre-sound baseline.

### Behavioral Electrophysiology Recordings

Prior to electrophysiology and behavior, positioned in the behavioral apparatus as they had been during training, and a 128-channel electrode (128D or 128M, Masmanidis Lab) was lowered into the secondary motor cortex orthogonal to the cortical surface (Yang et al., 2020). An optical fiber (ThorLabs, 400um, 0.39NA) was lowered on the surface of the exposed auditory cortex. The electrode was connected to a digitizing head stage (Intan) and electrode signals were acquired at 20khz, monitored in real time, and stored for offline analysis (OpenEphys). The probe was allowed to settle for at least 20 minutes, after which passive sound presentation and optical stimulation of auditory cortex for photo-tagging cells recorded in M2 was performed. For optical stimulation, the optical fiber was connected to a 470 nm laser source, calibrated to achieve 5-10 mW with 50 millisecond pulses of light. The duration of the laser pulse was programmatically controlled via TTL signals sent to the OptoEngine. Pulses were repeated 100 times with a randomized 0.5-1 second inter-pulse interval. Optical stimulation was followed by passive sound presentation, during which 10 frequencies of pure tones (0-32kHz) were presented 50 times each, randomly interleaved, with a 0.5-1 second inter-sound interval.

Following sound and laser stimulation, the lever was introduced and mice were allowed to make lever movements at will as in any other training session. In silent-trained mice, the novel tone was introduced at random, by the experimenter, after ~70-100 silent trials that had reached the movement goal. Auditory feedback was delivered as previously described for tone-trained mice. In all silent-trained mice, the novel tone was 12kHz, chosen because preliminary experiments indicated this frequency as the center of the average tuning curve for sound responsive cells. After the behavior was complete, the lever was removed and another session of optical stimulation and passive sound presentation was performed.

### Electrophysiology Data Analysis

After recording, electrical signals were processed and the action-potentials of individual neurons were sorted using Kilosort2, and manually reviewed in Phy2 based on reported contamination, stability of firing rate across the recording session, uniformity of the distribution of spike amplitudes, and inter-spike interval histograms. Peri-stimulus time histograms were created by analyzing a window of 500ms before and after an event (i.e. movement threshold, tone, etc). Smoothed average neural activity traces were made by taking the mean spiking activity across trials, and convolving this average with a hanning filter of 25-100 milliseconds width. All PSTHs shown were baseline-subtracted relative to activity during the 400 milliseconds window prior to the event.

For Z-score analyses, the 400 millisecond window before event onset was used to estimate the mean and standard deviation. Neurons were determined to be movement responsive if the firing rate in a baseline window (−900 to −500 milliseconds before the lever reached the movement-goal was significantly different than firing rates in any of three epochs during the movement: −200 to 0, 0 to 200, and 200 to 400 (all in milliseconds), using a Wilcoxon rank sum test (p<0.05). Neurons were determined to be sound responsive if the firing rate in the 200 millisecond window after tone onset differed from the 200 millisecond window before tone onset (Wilcoxon rank sum test, p<0.05) and if the latency to the peak response was less than 150 milliseconds.

To decode lever position from M2 data, we first identified every lever movement that reached the movement-goal. We then chunked neural activity into 100 millisecond bins and counted the number of action potentials produced by each neuron during the bin, as well as the average lever position during the bin. We then trained a linear decoder (support vector machine) with 80% of data and predicted the lever position from M2 activity with the held out data, repeated 100 times, each time with different held out data. We created confusion matrices by averaging the output of the linear decoder across all repeats.

To compare neural activity in silent-trained mice that are introduced to a self-generated sound, we compared neural activity on the last 30 trials of silent movements and the first 30 trials of sound-generating movements (0-200 milliseconds after reaching the movement-goal) (Wilcoxon rank sum test, p < 0.05). The slope of the relationship between passive tone and lever tone responses (Fig. 4D and 6J) was determined using a least squares fit to the 1st polynomial degree (MATLAB polyval and polyfit). R-values were calculated using the Pearson correlation coefficients (MATLAB corrcoef) between the two comparison groups, and p-values were calculated using a two-sided t-test. Tone and movement weights (Fig. 4) were determined with a multiple linear regression analysis (MATLAB regress), using a neuron’s passive tone response and response on silent movements as predictors of its response on sound-generating movements. Responsiveness to laser stimulation was determined by comparing cells’ firing rate in the 200 milliseconds window pre-laser onset versus the 200 milliseconds window post-laser onset (Wilcoxon rank sum test, p < 0.05).

In the comparison of AC-projecting M2 neurons’ activity on silent versus toned trials (Fig. 5H and I), the firing rate analysis window was restricted to 100-200 milliseconds. We performed a bootstrap analysis on the sound-producing trials, repeated 1000 times, and calculated the number of instances when the bootstrap mean exceeded the mean of the silent-trial data.

For PCA analyses (Fig. 7), z-scored population PSTHs of all movement and sound responsive cells were used for the singular value decomposition. PSTHs for silent, toned, and passive tone responses were concatenated along the time dimension, resulting in an input matrix of the following dimensions: row: neural population size; column: 3 x duration of each PSTH. The resulting data projected onto the first two PCs was then divided correspondingly by time segments. The angle between vectors in PC space was determined first by taking the dot product between the vector of the tone dimension with the vector determined by a single time point within a neural trajectory (relative to the zero origin). The dot product was then divided by the product of multiplying the magnitudes of the two initial vectors, and the inverse cosine taken of the resulting value. All resulting angles were converted to real numbers. Errorbars on angle and projection plots were calculated using a bootstrap analysis, where PCA was repeated 10,000 times with populations whose cell identities were resampled with replacement. Angles and projections were recalculated on each repetition and the standard deviation taken across all repetitions.

To estimate the relative influence of tone frequency and startle movements on sound-evoked responses in M2, we fit a linear regression model that included the startle magnitude on each trial, the sound frequency on each trial, and interaction terms between startle magnitude and frequency (MATLAB fitlm). We then estimated how each term contributed to the population-level firing of M2 neurons on each trial of sound playback. We report the T-statisic for each term in the model, averaged across mice.

### Histology and Image Quantification

Tracing of connections from auditory cortex (AC) to secondary motor cortex (M2) were performed with injections of anterogradely-delivered Td-Tomato in AC (AAV-CAG-TdTomato, UNC, titer 5.3e12) and retrogradely-delivered GFP in M2 (retroAAV-hSyn-EGFP, Addgene, titer 4.2e12). M2 was injected (1.0-1.5mm AP, 0.5-0.7mm ML) at three depths (200um, 400um, and 600um), 100nL at each depth, for a total injection volume of 300nL. AC was injected (−2.5mm AP, ~4.2mm ML) at a ~40 degree angle, nearly orthogonal to the cortical surface with the same depths and injection volume as M2. Viruses were allowed to express for two weeks, after which perfusions were performed and the brain refrigerated in formaldehyde. For processing, brains were encased in agar and sliced on a vibratome in 70um slices. Frontal and temporal cortical sections were kept and stored in PBS. Terminals of anterograde tracings (both TdTomato and ChR2), were blocked in 5% Normal Goat Serum (Rockland, Abcam) in 0.1% PBS TritonX, then amplified with a primary incubation in Rabbit-anti-RFP (Rockland, 1:1000 dilution in PBS) and secondary incubation in Goat Anti-Rabbit Alexa 594 (Abcam, 1:2000 dilution in PBS). Cell bodies and injection sites were not amplified. All sections were mounted on slides and imaged under a Leica confocal microscope. AC terminal fluorescence was analyzed using ImageJ, while AC cell bodies were manually counted. Boundaries between cortical regions were identified using the Franklin and Paxinos (2007) brain atlas, and cell layers were estimated with DAPI-stained images and the Allen Brain Atlas.

### Muscimol Silencing and Analysis

During muscimol silencing experiments and saline control experiments, a small piece of gelfoam soaked in saline was placed over auditory cortex. After 500 presentations of passive sounds, the gelfoam was removed and replaced with gelfoam soaked in room-temperature muscimol (0.5 mg/mL). We presented another 500 sounds immediately after muscimol placement, waited 30-45 minutes, and presented another 500 sounds. We compared neural responses presented during saline application and during the final window of muscimol application.

### Statistical Analysis

All sizes of animal cohorts are denoted by a capital N, while cell population sizes are denoted by a lowercase n. Unless otherwise noted, all errorbars represent standard error (standard deviation divided by square root of the size of the analysis population). A two sided, non-parametric Wilcoxon rank sum test (MATLAB ranksum), was used for most statistical comparisons including significant responsiveness to stimuli (i.e. sound, laser, etc.) and comparisons between differing subpopulations or training cohorts. For comparison within and between tone and movement responsive subpopulations (Supplemental Fig. 4C,4D), a parametric paired t-test (MATLAB ttest) and two-sided t-test (MATLAB ttest2) were used, respectively. Statistical comparisons of the activity of simultaneously recorded cells between conditions were tested for normality using a Kolmogorov-Smirnov test (MATLAB kstest). If normal, we then used a paired t-test (MATLAB ttest) for significance and if not normal, a Wilcoxon signed rank test (MATLAB signrank) was used. Statistical significance on figures was denoted as *p<0.05, **p < 0.005, ***p<.0005

## Acknowledgements

We thank Dr. Nick Audette, Dr. Alessandro La Chioma, and Ariadna Corredera for their feedback on this manuscript. We thank Drs. Kathy Nagel, Christine Constantinople, and Dan Sanes, as well as members of the Schneider lab for their input throughout the project. We thank WenXi Zhou for guidance on analyzing population dynamics, and Dr. Alessandro La Chioma for his input on their interpretation. We thank Hoda Ansari and Jessica A. Guevara for their assistance with managerial support and animal care. This research was supported by the National Institutes of Health (T32-MH19524-27 to B.E.H., 1F31-DC019307 to B.E.H., 1R01-DC018802 to D.M.S.); a Career Award at the Scientific Interface from the Burroughs Wellcome Fund (D.M.S); fellowships from the Searle Scholars Program, the Alfred P. Sloan Foundation, and the McKnight Foundation (D.M.S.). D.M.S. is a New York Stem Cell Foundation - Robertson Neuroscience Investigator.

